# Characteristics of human encounters and social mixing patterns relevant to infectious diseases spread by close contact: A survey in southwest Uganda

**DOI:** 10.1101/121665

**Authors:** O le Polain de Waroux, S Cohuet, D Ndazima, A J Kucharski, A Juan-Giner, S Flasche, E Tumwesigye, R. Arinaitwe, J Mwanga-Amumpaire, Y Boum, F Nackers, F Checchi, R F Grais, W J Edmunds

## Abstract

Quantification of human interactions relevant to infectious disease transmission through social contact is central to predict disease dynamics, yet data from low-resource settings remain scarce. We undertook a social contact survey in rural Uganda, whereby participants were asked to recall details about the frequency, type, and socio-demographic characteristics of any conversational encounter that lasted for ≥5 minutes (henceforth defined as ‘contacts’) during the previous day. An estimate of the number of ‘casual contacts’ (i.e. <5 minutes) was also obtained. A total of 568 individuals were included. On average participants reported having routine contact with 7.2 individuals (range 1-25). Children aged 5-14 years had the highest frequency of contacts and the elderly (≥65 years) the fewest (P<0.001). A strong age-assortative pattern was seen, particularly outside the household and increasingly so for contacts occurring further away from home. Adults aged 25-64 years tended to travel more and further than others, and males travelled more frequently than females. Our study provides detailed information on contact patterns and their spatial characteristics in an African setting. It therefore fills an important knowledge gap that will help more accurately predict transmission dynamics and the impact of control strategies in such areas.

## INTRODUCTION

Quantification of human interactions relevant to the spread of these infectious diseases is essential to accurately predict their infection dynamics and the impact of control strategies [1, 2].

Detailed surveys of social mixing patterns have now been undertaken in a number of settings [2-13]. Studies have shown that people tend to mix with other individuals of their own age (i.e. assortative mixing); however, the frequency of contact, the degree of intergenerational mixing and the characteristics of mixing tend to vary between settings, depending on factors such as household size, population density and local activities, among others [3-11].

Data from low-resource settings remain scarce, with only three studies in Africa published to date [10, 12, 13], and none from Uganda.

With the exception of a recent study from China [11], the spatial dispersal of social contacts relevant for transmission has often been overlooked, and there is – to our knowledge – no published information from low-income settings. Spatial mobility is particularly important for epidemic risk prediction of novel and re-emergent diseases, and for the optimization of routine control programmes [14].

To address this knowledge gap, we set up a study of social contacts relevant to the spread of infections transmitted through the respiratory route or by close contact, in rural southwest Uganda.

## MATERIALS AND METHODS

The study was conducted in four sub-counties of Sheema North Sub-District (southwest Uganda), an area with a total of about 80,000 inhabitants. About half (49%) of the district’s population is <15 years. The area is primarily rural.

### Study design

We conducted a two-stage age-stratified community-based study in Sheema North Sub-district between January and March 2014 on a subset of individuals included in a population-based survey of nasopharyngeal carriage of *Streptococcus pneumoniae* (Nackers F et al. manuscript in preparation). The sample size calculations by age group for the nasopharyngeal carriage study included 538 children <2 years, 323 children aged 2 – 4 years, 583 aged 5 – 14 years and 327 individuals aged ≥15 years. Using the same age groups, but different inclusion probabilities, we estimated that, based on previous results [12, 15], including all 327 individuals ≥15 years and a subset of 90 children <2 years, 90 children aged 2-4 years and 180 children aged 5 – 14 years, for a total target sample size of 687, would provide a precision of just over 1 contact on the mean number of contacts per day, and enable detection of a 20% difference in the average number of daily contacts by age group, accounting for 10% non-response.

Individuals were selected from 60 clusters randomly sampled from the exhaustive list of 215 villages and two small towns in the sub-county, with an inclusion probability proportional to the size of the village or town. Within each cluster 11 or 12 households were randomly selected and in each household only one individual was selected for inclusion in the study. A household was defined as a group of individuals living under the same roof and sharing the same kitchen on a daily basis. One individual from each household was randomly selected from within a predefined age group based on a random sequence of age groups according to the age group sampling quota by cluster.

When nobody in the household was from that age group, either someone from another age group was selected providing that the quota for that age group had not been reached in the cluster, or the closest neighbouring household was visited instead. In case of non-response, another attempt was made later in the day or the following Saturday. Survey teams had a day off on Thursdays and Sundays.

### Data collection

Informed consent was sought for individuals aged > 13 years, and consent was sought from a parent or guardian otherwise.

Ethnical approval was obtained from the Ethical review boards of Médecins Sans Frontières (MSF), the Faculty of Medicine Research & Ethics Committee of the Mbarara University of Science and Technology (MUST), the Institutional Ethical Review Board of the MUST, the Uganda National Council for Science and Technology (UNCST) and the London School of Hygiene and Tropical Medicine (LSHTM).

Participants were asked to recall information on the frequency, type and duration of social encounters from the time they woke up the day before the survey until when they woke up on the survey day (˜ 24 hours).

We defined contacts as individuals with whom there was at least one two-way conversational encounter (three or more words) lasting for ≥5 minutes. Participants were first asked to list all the places they had visited in the previous 24 hours, the number of people they had contact with, their relationship with each individual mentioned, the age (or estimated age) of each listed contact and how long the encounter lasted for. Contacts involving skin-to-skin touch or sharing utensils passed directly from mouth-to-mouth were defined as ‘physical’ contacts.

We defined as ‘casual contacts’ short conversational encounters lasting less than 5 minutes. Participants were only asked to estimate the number of casual contacts they had, based on pre-defined categories (<10, 10-19, 20-29, ≥30), but were not asked to provide detailed information about the nature of the encounter or the socio-demographic characteristics of the person met. Casual contacts are generally inaccurately reported in social contact surveys [7], particularly in a retrospective design, and most contacts important for the transmission of respiratory infections are believed to be close rather than casual [6].

The questionnaire was designed in English, translated to Ruyankole, the local language, and back-translated to English for consistency (Supporting Information Text S1). For children <5 years, parents were asked about their child’s encounters and whereabouts. Children aged 5 – 14 years were interviewed directly, using an age-appropriate questionnaire.

Geographical coordinates from each participant’s household and of the centre point of each village were taken using handheld GPS devices.

Questionnaires completed in the field were double entered on a preformatted data entry tool by two data managers working independently. Data entry conflicts were identified automatically and resolved as the data entry progressed.

### Analysis

#### Characteristics of social contacts by time, person and place

We analysed the frequency distribution of contacts for a set of covariates, including age, sex, and occupation, day of the week, distance travelled, and type of contact. Encounters reported with the same individual more than once counted as one contact only. Distance travelled was measured as straight line distances between the centre point of the participants’ home village/town and that of the village/town where each reported encounter took place.

We used negative binomial regression to estimate the ratio of the mean contacts per person as a function of the different covariates of interest. Negative binomial was preferred over Poisson regression given evidence of over-dispersion (variance > mean, and likelihood ratio significant (P<0.05) for the over-dispersion parameter). We considered variables associated with contact frequency at p<0.10 for multivariable analysis, and retained them in multivariable models if they resulted in a reduction of the Bayesian Information Criterion (BIC).

Next, we explored whether people reporting a high frequency of casual contacts (≥10 casual contacts) differed from those reporting fewer contacts with regards to their socio-demographic characteristics. We did so using log-binomial regression to compute crude and adjusted relative risks (RRs) for having a high frequency In all analyses we accounted for possible within-cluster correlation by using linearized based variance estimators [16]. Analyses were also weighted for the unequal probabilities of sampling selection by age group.

#### Age-specific social contact patterns

We analysed the age-specific contact patterns through matrices of the mean number of contacts between participants of age group *j* and individuals in age group *i*, adjusting for reciprocity, as in Melegaro et al. [6].

If *X_ij_* denotes the total number of contacts in age group *i* reported by individuals in age group *j*, the mean number of reported contacts (*m_ij_*) is calculated as *x_ij_*/*P_j_*, where *P_j_* is the study population size of age group *j*. At the population level the frequency of contacts made between age groups should be equal such that *m_ij_P_j_* = *m_ji_P_i_*. The expected number of contacts between the two groups is therefore *C_ij_* = (*m_ij_P_j_* + *m_ji_P_i_*)/2. Hence, the mean number of contacts corrected for reciprocity (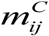) can be expressed as *C_ij_*/*P_j_*.

#### Epidemic simulations

Finally, in order to explore the infection transmission dynamics resulting from our contact pattern data, we simulated the spread of an immunizing respiratory infection transmitted through close contact in a totally susceptible population. The model contained nine mixing age groups, with a transmission rate at which individuals in age group *j* come into routine contact with individuals in age group *i* computed as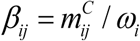, where *ω_i_* is the proportion of individuals in age group *i*, and 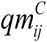 is the next generation matrix, with *q* representing the probability of successful transmission per contact event [17]. We assumed *q* to be homogeneous and constant across all age groups and conducted a set of simulations for fixed values of *q* between 25% and 40%, in line with what has been reported with influenza pandemic strains [17, 18]. The basic reproduction number (*R*_0_) – which corresponds to the average number of people infected by one infectious individual in a totally susceptible population – was calculated as the dominant eigenvalue of the next generation matrix. We took uncertainty estimates in the contact matrices (and hence final size outputs) into account by iterating the model on bootstrapped matrices.

We then computed the final epidemic size (i.e. the number of individuals who would have been infected during the epidemic) for each specific age group, based on a mass action model adapted to account for multiple age classes, as described in Kucharski et al. [19].

Estimates obtained using the contact data from Uganda were compared to that of Great Britain, using data from the POLYMOD study [4] for the latter and a similar approach to compute the mixing matrix. The model was parameterised with social contact data on physical contacts only, lasting ≥5 minutes, rather than all contacts, given that physical contacts generally seem to better capture contact structures relevant for the transmission of respiratory infections [6], and that the definition of physical contacts is more similar and comparable between studies than that of overall contacts.

All analyses were performed in STATA 13.1 IC and R version 3.2.

## RESULTS

### Study population

A total of 568 individuals participated in the survey, but no information about age and contacts was missing for 2 individuals, resulting in 566 included in the analysis. This corresponds to an overall response rate of 83%; higher among ≥15 years old (98%), and lower among under 2s (68%), 2-4 year olds (64%), 5 –14y olds (69%).

There were more female (58%) than male respondents, but this differed by age group, with fewer females in young age groups and more adult females than males (Table S1 in the Supporting Information).

The mean household size was 5.3 (median 5, range 1 – 18). Almost all (98%) school-aged children aged 6 – 14 years attended school or college. Among adults, agriculture was the main occupation and about 27% of the females were homemakers/housewives (Table 1).

**Table 1.** Mean Number of Reported Contacts and Ratio of Means By Socio-demographic Characteristic of The Study Population, Sheema, Uganda, January – March 2014.

| Variables | N | Mean number of<br>contacts (95%CI) | Crude RoM<br>(95%CI) | Age adjusted RoM (95%CI) |
| --- | --- | --- | --- | --- |
| Age groups |  |  |  |  |
| <2y | 61 | 6.11 (5.42 ,6.81) | 0.99 (0.83 ,1.17) |  |
| 2-4y | 57 | 6.70 (6.08 ,6.81) | 1.08 (0.95 ,1.23) |  |
| 5-9y | 74 | 8.50 (7.79 ,9.20) | 1.37 (1.18 ,1.59) |  |
| 10-14y | 51 | 8.70 (7.52 ,9.90) | 1.40 (1.19 ,1.66) |  |
| 15-24y | 91 | 6.20 (5.51 ,6.88) | ref |  |
| 25-34y | 55 | 6.89 (5.99 ,7.80) | 1.11 (0.93 ,1.33) |  |
| 35-44y | 54 | 7.73 (6.99 ,8.46) | 1.25 (1.09 ,1.43) |  |
| 45-54y | 46 | 7.74 (6.37 ,9.11) | 1.25 (1.01 ,1.54) |  |
| 55-64y | 26 | 6.27 (5.01 ,7.53) | 1.01 (0.80 ,1.29) |  |
| 65+y | 48 | 4.85 (4.18 ,5.53) | 0.78 (0.64 ,0.95) |  |
| Sex |  |  |  |  |
| Female | 328 | 7.05 (6.66 ,7.44) | ref |  |
| Male | 235 | 7.47 (6.82 ,8.11) | 1.06 (0.96 ,1.18) |  |
| Occupation/daily activity |  |  |  |  |
| Pre-school child | 93 | 7.00 (6.30 ,7.70) | 1.06 (0.90 ,1.23) | 1.26 (1.02 ,1.55) |
| Student | 166 | 8.27 (7.56 ,8.98) | 1.27 (1.10 ,1.46) | 1.30 (1.05 ,1.63) |
| Office worker | 4 | 11.34 (9.64 ,13.03) | 1.81 (1.20 ,2.73) | 1.70 (1.32 ,2.18) |
| Shop worker | 34 | 6.82 (5.56 ,8.08) | 1.03 (0.84 ,1.25) | 1.03 (0.83 ,1.29) |
| Agriculture | 105 | 7.30 (6.58 ,8.01) | 1.12 (0.97 ,1.30) | 1.12 (0.96 ,1.30) |
| Other manual worker | 40 | 5.37 (4.36 ,6.38) | 0.85 (0.70 ,1.04) | 0.85 (0.68 ,1.06) |
| At home | 60 | 6.43 (5.62 ,7.24) | ref | ref |
| Unemployed | 11 | 6.44 (2.92 ,9.96) | 0.83 (0.60 ,1.15) | 1.22 (0.77 ,1.94) |
| Retired | 8 | 4.77 (3.75 ,5.80) | 0.71 (0.48 ,1.05) | 0.89 (0.70 ,1.13) |
| Other/unreported | 41 | 6.65 (5.80 ,7.51) | 1.02 (0.85 ,1.23) | 1.18 (0.98 ,1.43) |
| Day of the week |  |  |  |  |
| Weekday | 439 | 7.14 (6.79 ,7.49) | ref |  |
| Sunday | 124 | 7.50 (6.56 ,8.44) | 1.05 (0.92 ,1.20) | 1.04 (0.92 ,1.18) |
| Travel outside village/town in previous 24 hours |  |  |  |  |
| No | 427 | 6.57 (6.20 ,6.92) | ref |  |
| Yes | 139 | 9.04 (8.35 ,9.73) | 1.38 (1.25 ,1.52) | 1.35 (1.22 ,1.49) |
| Number of casual contacts |  |  |  |  |
| <10 | 315 | 5 (1 -15) | Ref | Ref |
| 10 -19 | 119 | 8 (2-23) | 1.43 (1.28, 1.59) | 1.39 (1.25, 1.55) |
| ≥20 | 56 | 9 (2-25) | 1.64 (1.44; 1.86) | 1.61 (1.43,1.83) |
| Don't know | 76 | 8 (0-19) | 1.52 (1.37; 1.68) | 1.45 (1.29, 1.64) |

Footnote CI=Confidence Interval; RoM: Ratio of Means

### Characteristics of contacts

#### Contacts (i.e. ≥5 minutes long)

A total of 3,965 contacts with different individuals were reported, corresponding to an average of 7.2 contacts per person (median 7, range 0 - 25) (Figure 1). The majority of contacts were physical (mean 5.1, median 5 (range 0 – 18)).

**Figure 1:**
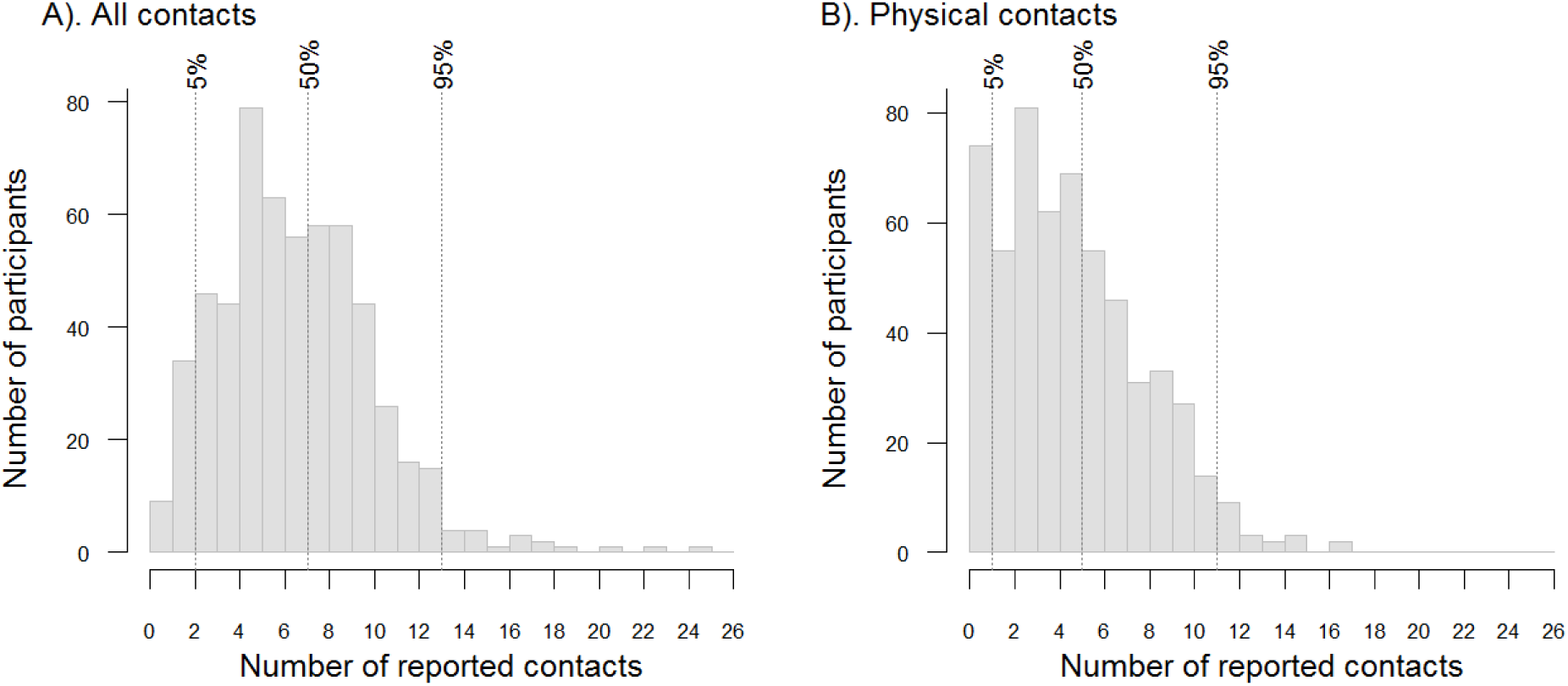
Number of Reported Contacts, Including All Contacts (A) and Physical contacts (B), Sheema, Uganda, January – March 2014. <u>Legend:</u> the vertical dotted lines represent the 5% centile, the median and 95% centile of the total number of reported contacts

Over half of all contacts (n= 2,060 (52%)) were with household members, 627 (16%) with other relatives, 873 (22%) with colleagues/friends/schoolmates and 402 (10%) with other individuals. The duration of contacts is shown in Figure S2 (Supporting Information).

Most contacts (82%) were with individuals who would be normally seen daily, 520 (13%) with people normally seen at least weekly, 4% with people met more rarely and 1% of the reported contacts were with people that the participants had never met before.

We found marked differences in the number of contacts by age group, but not by sex. School-aged children reported the highest daily number of contacts, while the elderly had the fewest (Table 1). Table 1 provides further details about the population characteristics, the mean number of contacts by socio-demographic and other covariates, as well as the ratio of mean contacts by covariate. Results were adjusted for age, but not other variables, as identified through the results of the negative binomial model.

Overall, contacts tended to be assortative, as shown by the strong diagonal feature on Figure 2, with most of the intergenerational mixing occurring within households (Figure 3). Only teenagers and adults reported non-physical contacts (Figure 3). Reciprocity correction accounted for the differential reporting between age groups, particularly higher frequency of contacts reported by small children with older age groups than older age groups reported (Figure S2 in Supporting Information).

**Figure 2:**
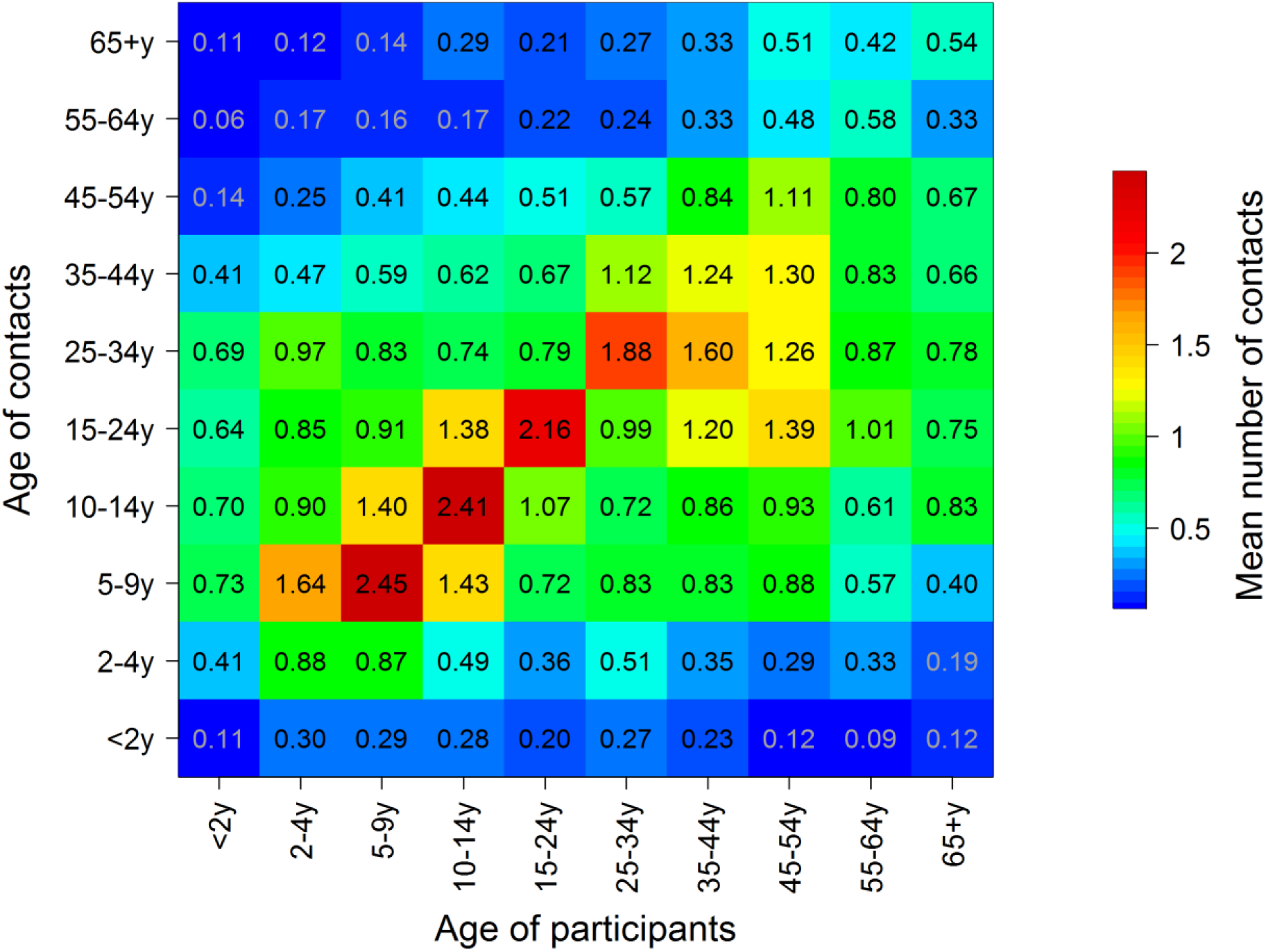
Average Number of Reported Contacts By Age Group, Sheema, Uganda, January – March 2014 <u>Legend:</u> Numbers in each cell represent the average number of contacts between between age groups corrected for reciprocity, and 95% confidence intervals are shown

**Figure 3:**
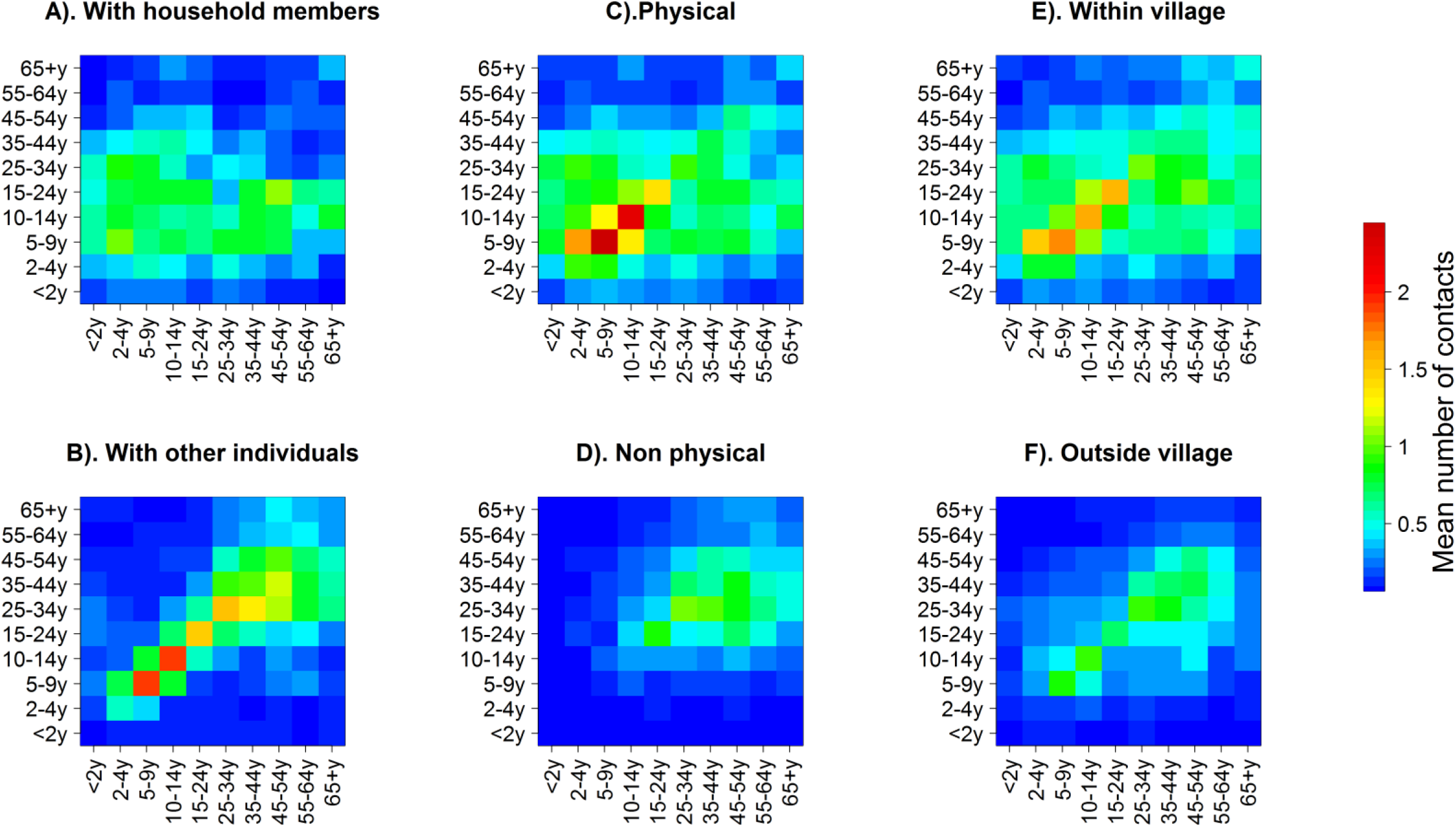
Contact Matrices With Household members and Non-Household Members (Left upper and lower panel), for Physical and Non-Physical Contacts (Middle upper and lower panel), and for Contacts Made Within and Outside the Village (Right upper and lower panel), Sheema, Uganda, January – March 2014

There was no statistical difference in the average number of contacts between weekend (Sunday) and weekdays (Monday, Tuesday, Thursday and Friday) (Table 1). Given that survey teams had a day off on Sundays and on Thursdays information about contact on Saturdays and Wednesdays was not recorded. About a quarter (n=136 (24%)) of participants reported social encounters outside their village of residence, and about 12% of contacts occurred outside participants’ village of residence. The majority (56%) of people who travelled outside their village went to places located within a 5km radius from the centre point of their village of residence, and 90% stayed within 12km (Figure 4). Adult males tended to travel more than females (Figure 4). Most contacts made outside the household as well as those with individuals outside participants’ village were mostly assortative (Figure 3), and the proportion of contacts outside the village was different by age group (P<0.001); higher among adults, increasingly so as distance from home increased (Figure 4).

**Figure 4:**
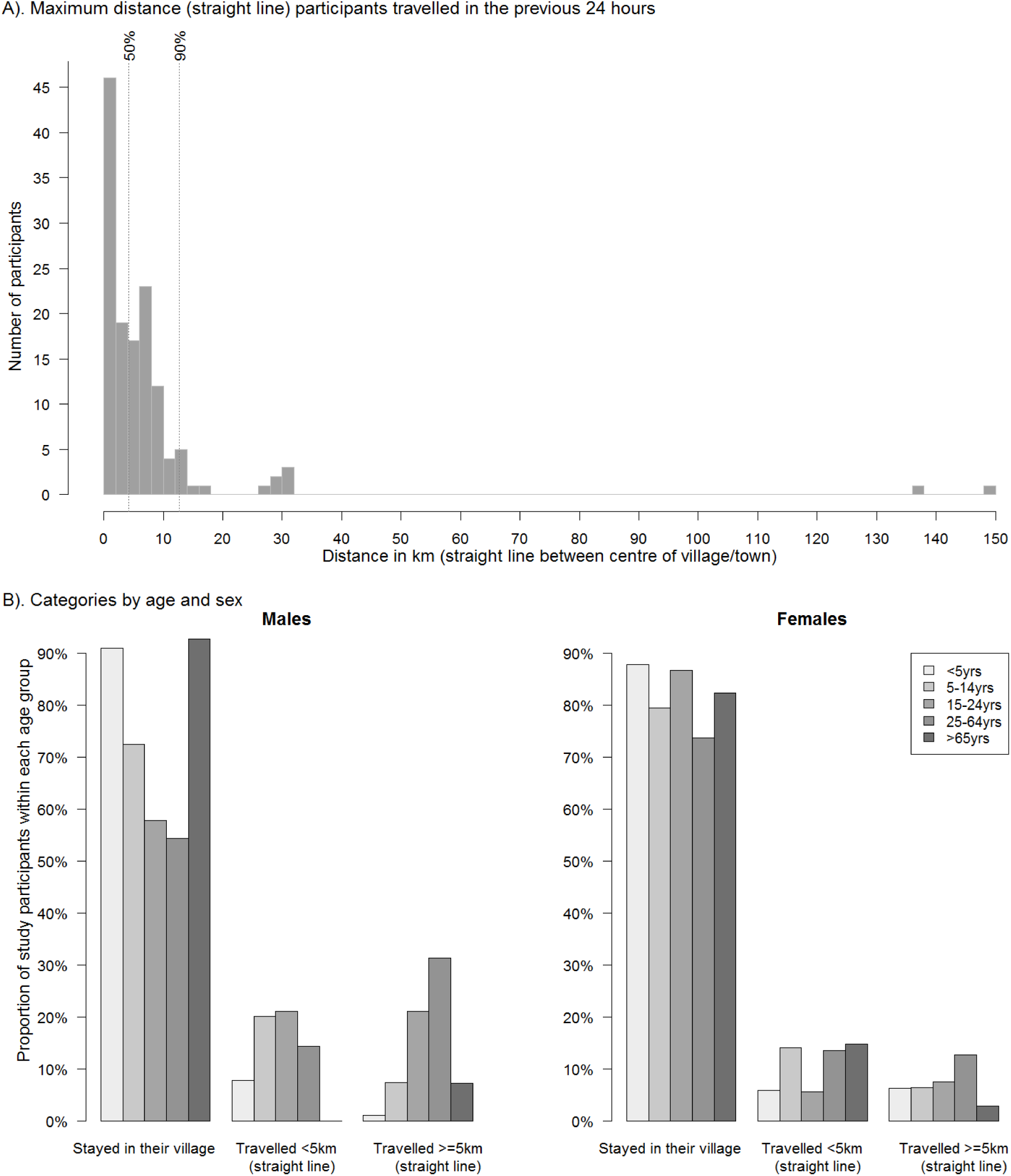
Distance Travelled By Study Participants in the 24 Hours Preceding the Survey, Overall (A) and By Categories of Distance, Age and Sex (B), Sheema, Uganda, January – March 2014

#### ‘Casual’ contacts (<5 minutes long)

Information on the number of casual contacts was reported by 490 (87%) participants. Among those, 64% (n=315) estimated they had fewer than 10 different contacts, 24% reported between 10 and 19 casual contacts, 6% reported between 20 – 29 contacts and 6% reported an estimated 30 contacts or more.

Individuals who reported high levels (i.e. ≥10 contacts) of social contacts also tended to report more contacts (Table 1). We found no difference between those reporting high number of social contacts (≥10) and others, by age, sex or day of the week (Table S2 in the Supporting Information). However, people whose primary activity was at home tended to reported fewer casual contacts than others, and there were about 50% more individuals reporting high levels of casual contacts among those who travelled outside their village.

#### Epidemic simulations

Finally, we compared patterns of reported physical contacts in Uganda and Great Britain, and explored differences in the relative and absolute epidemic size by age group, as well as the corresponding *R*_0_, for a hypothetical respiratory infection in an immune-naive population.

The number of reported physical contacts was similar between Uganda and Great Britain, with the average number of contacts by age group ranging from 3.2 (≥65 year olds) to 7.3 (2 – 4 year olds) in Uganda and from 3.3 (55 – 64 year olds) to 7.3 (10 – 14 year olds) in Great Britain. However contacts were more assortative in Britain than in Uganda (Figures 5A & B), some of which might be related to differences in household structures and number of household contacts, as contacts outside the household were mostly assortative (Figure 3).

The computed mean values of *R*_0_ for a per contact infectivity value (*q*) ranging from 0.25 to 0.40 was slightly higher in Great Britain than in Uganda (1.51 to 2.41 vs. 1.40 to 2.24). Figure 5F shows the values for an infectivity parameter of 0.33. The proportion of people infected in younger age groups was also higher in Great Britain, and there were proportionally more adults infected in Uganda. However, given the differences in population structure, the total number of infections in the population was higher in Uganda than in Great Britain (Figures 5 C – E).

**Figure 5:**
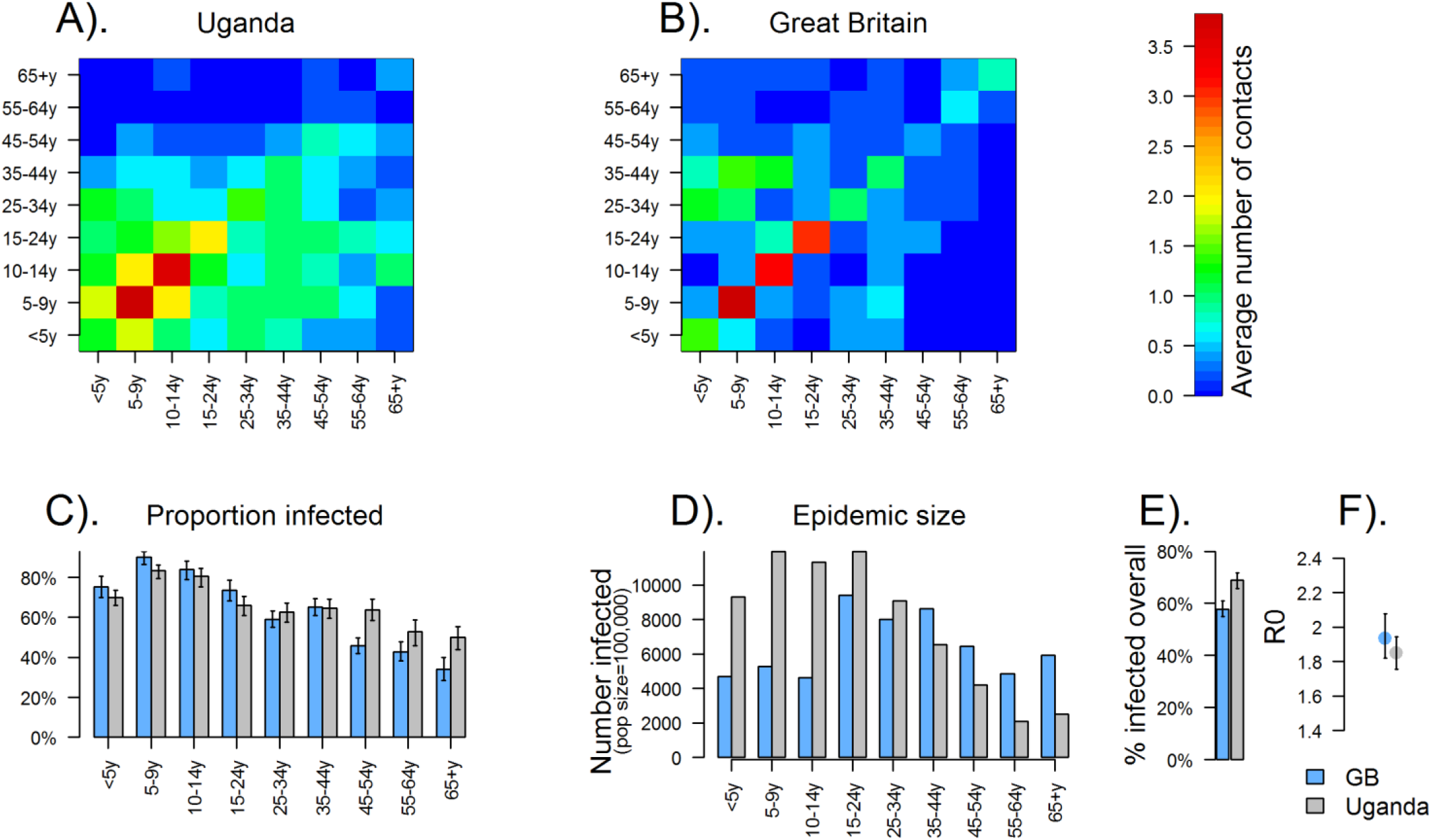
Epidemic Simulation Using Matrices on Physical Contacts from Uganda (A) and Great Britain (B), for a Hypothetical Respiratory Infection In An Immune-Naïve Population, with the Proportion Infected by Age Group (C), the Epidemic Size by Age group (D), the Overall Proportion Infected (D) and the Basic Reproduction Number R0 (F). <u>Legend:</u> A:Matrix for physical contacts in Uganda. B: Matrix of physical contacts in Great Britain. C: Epidemic final size simulation: Proportion of individuals infected by age group in Great Britain (blue) and Uganda (grey), with error bars representing the 95% confidence interval. The results are presented for a *q* value of 33%. D: Epidemic size by age group, based on a total population size of 100,000 in Great Britain and in Uganda. E: Total proportion of people who were infected at the end of the epidemic in each setting. F: Estimates of *R*_0_ for each setting, based on a *q* value of 33%, with dots showing the mean value and the bars showing the 95% CI

## DISCUSSION

To our knowledge this is only the third study of its kind in Africa [10, 12], and the first one to specifically explore spatial patterns of social contacts. The quantification of mixing patterns is central to accurately model transmission dynamics and inform infectious disease control strategies [4]. Having such data thus fills an important gap, particularly given the high burden of respiratory infections in low income settings [20, 21], and the risk of emerging and re-emerging diseases transmitted by close interpersonal contact, such as influenza [22], measles [23] or meningitis [24].

Our findings share similarities with studies from Africa [10, 12, 13] and other low or lower-middle income settings [15, 25], including the high contact frequency among school-aged children and that most contacts tend to be age-assortative. We also found substantial mixing between age groups, largely driven by intra-household mixing. This may result in a higher force of infection from children to adults than would be seen in other contexts such as Great Britain, as our final size epidemic model suggests. The final size model should be seen as an illustration of how different social mixing patterns impact on disease epidemiology in different settings, rather than a specific quantification of the differences. It shows the importance of using setting-specific data when modelling disease dynamics and evaluates control strategies. Our data could be best applied to evaluate transmission dynamics and the impact of interventions for endemic diseases and current epidemics in non-naïve population in similar rural East African contexts. In our final size model, it is also likely that our retrospective design resulted in underreporting compared to a prospective diary-based approach [26], which hampers comparisons between countries. In sensitivity analyses we explored the impact of potential underreporting in our retrospective survey design compared to a prospective diary-based approach [26], assuming a 25% under-ascertainment compared to a diary-based study, with homogeneous underreporting across age groups. In such scenario, the proportion of infections across all age groups is predicted to be higher in Uganda than in Britain, disproportionally so in adults, and the *R*_0_ to be higher too (see Figure S3 in the Supporting Material).

Our results also provide important insights into the local spatial dynamics of routine daily human interactions, showing that most contacts tend to occur within the vicinity of people’s area of residence, that working age adult males travel most and young children and the elderly the least, and that contacts tend to be increasingly age assortative as people travel further away from home. Similar patterns were observed in rural and semi-urban China [11]. Such findings have important implications to predict outbreak dynamics and control strategies given that interconnectedness between geographic patches is an essential factor driving epidemic extinction or persistence of epidemics hotspots and the effectiveness of control strategies. Studies of measles in Niger suggest that dynamics differ from that observed in high-income countries in the pre-vaccination era, likely due to different mixing patterns and weaker spatial connectivity [27, 28]. This, together with important variations in vaccination coverage between local geographic patches [29-31], strengthens the need to account for spatial mobility when designing efficient control strategies in those settings. Optimal targeted interventions tailored to specific geographic clusters of high transmission have also been key considerations in recent cholera outbreaks in Africa, given the limited available vaccine doses [32, 33]. Spatially targeted approaches are also central to outbreak control in the recent West African Ebola epidemic [34], and the current measles epidemic in the Democratic Republic of the Congo, which is sustained in part due to

In our study the frequency of contacts was about half that of the number of contacts reported in Kenya, [10] or South Africa [12], and lower than in a recent contact study conducted by Melegaro et al. in rural and peri-urban areas in Zimbabwe [13]. Although differences between settings are expected, some of these are likely to be due to the exclusion of ‘casual contacts’ from our contact count. There might be further differences linked to the definition of social contacts, which was based on conversational encounters in our study but not in the Kenyan study [10]. When defining contacts based on conversational exchanges the household setting tends to dominate over other settings, compared to a more inclusive definition [8].

Both our contact definition and the retrospective study design may have also resulted in some level of reporting bias with more stable, regular contacts being reported over others. However, the extent to which a more inclusive definition reflects contact events relevant for transmission remains unclear. Modelling studies suggest that close interpersonal rather than short casual contacts matter more for transmission of respiratory infections [6]. In addition, for modelling purposes the age-specific structure of relative contact frequency matters more than the actual reported frequency, as matrices are scaled to fit epidemiological data. Our retrospective interview-based design thus offers a simpler and easier alternative to prospective diary based approaches, particularly in such setting. Further research should explore what contact information is most relevant and how such data should best be captured.

Selection bias may have occurred to some extent, particularly given that more adult women were included than men. However, there was no significant difference in the number of contacts reported between males and females, including at the weekend, suggesting that selection bias was unlikely to be major. We also tried to reduce selection bias by interviewing on Saturdays people who were initially absent on the survey.

In conclusion, our study fills an important gap for two main reasons. First, we provide information by detailed age groups about social contacts and mixing patterns relevant to the spread of infectious diseases in a region where such data are scarce. Second, we also provide some insights into spatial characteristics of social encounters. Although this has increasingly being recognized as an important component in evaluating epidemic risk and in the design of efficient control strategies, it has not previously been quantified in low-income settings, and should be explored further. Our study thus provides essential evidence to inform further research and infectious disease modelling work, particularly in similar rural African settings.

## FUNDING

This work was supported by Médecins Sans Frontières International Office, Geneva, Switzerland. Epicentre received core funding from Médecins Sans Frontières. OLP was supported by the AXA Research Fund through a Doctoral research fellowship. The funders had no role in the study design, data collection, data analysis, data interpretation or write up of the manuscript.

## ACKNOWLEDGMENTS

We would like to thank all the surveyors for their work. We are grateful to Gertrude Ngabirano for her help in translating questionnaires.

## CONFLICT OF INTEREST

The authors declare they have no conflict of interest.

## Web material

### Tables

**Table S1.** Age and Sex Distribution of Study Participants

| Age category | Number (%) female | Number (%) male | Total number |
| --- | --- | --- | --- |
| <2 years | 23 (38%) | 38 (62%) | 61 |
| 2 – 4 years | 26 (45%) | 31 (55%) | 57 |
| 5 – 9 years | 37 (50%) | 37 (50%) | 74 |
| 10 – 14 years | 22 (43%) | 29 (57%) | 51 |
| 15 - 24 years | 53 (58%) | 38 (42%) | 91 |
| 25 – 34 years | 45 (79%) | 12 (21%) | 57 |
| 35 – 44 years | 35 (64%) | 20 (36%) | 55 |
| 45 – 54 years | 35 (76%) | 11 (24%) | 46 |
| 55 – 64 years | 20 (77%) | 6 (23%) | 26 |
| 65+ years | 34 (71%) | 14 (29%) | 48 |

**Table S2.**
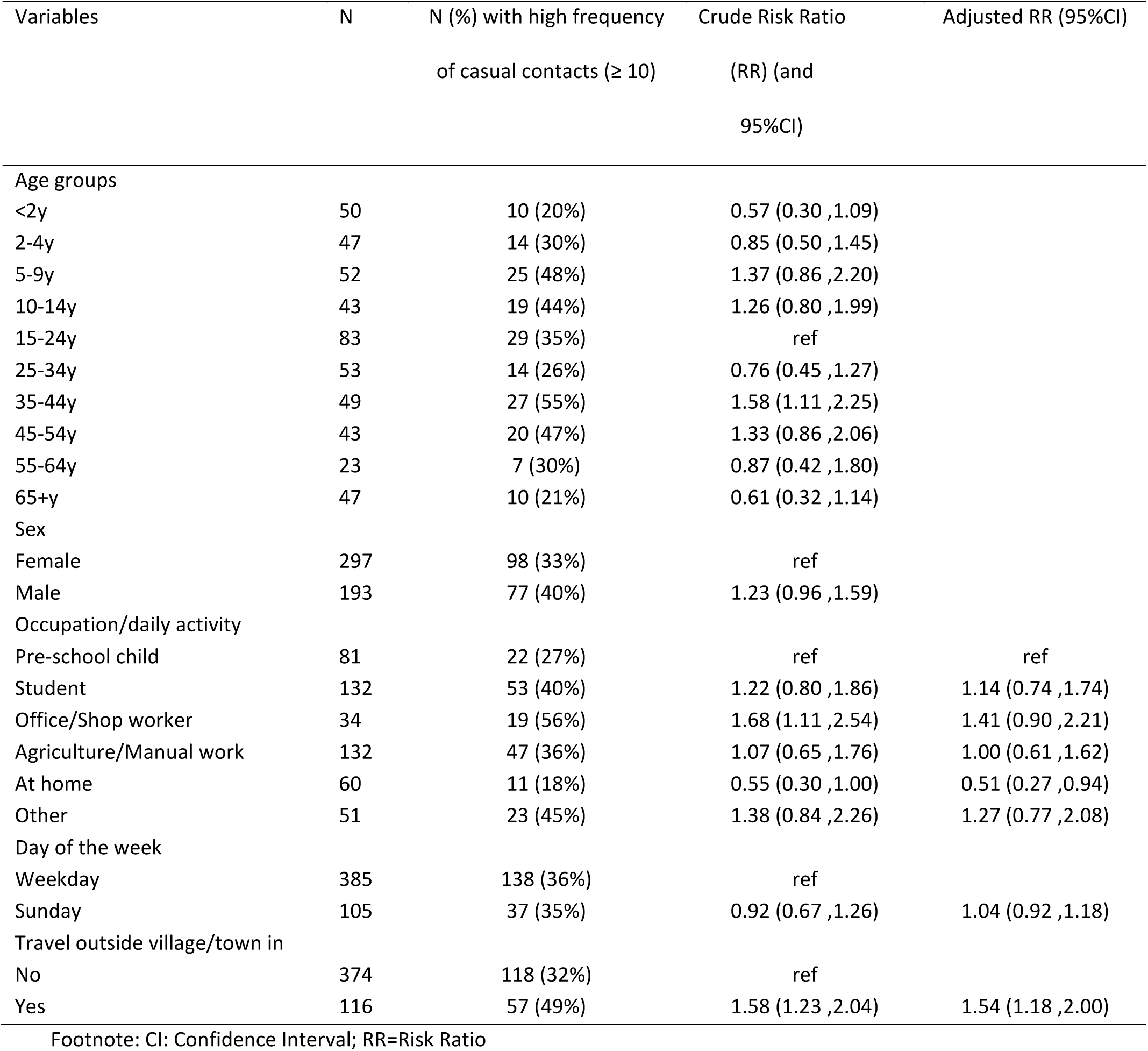
Association Between Socio-demographic Variables and Level of Social Contacts

### Supplementary figures

**Figure S1.**
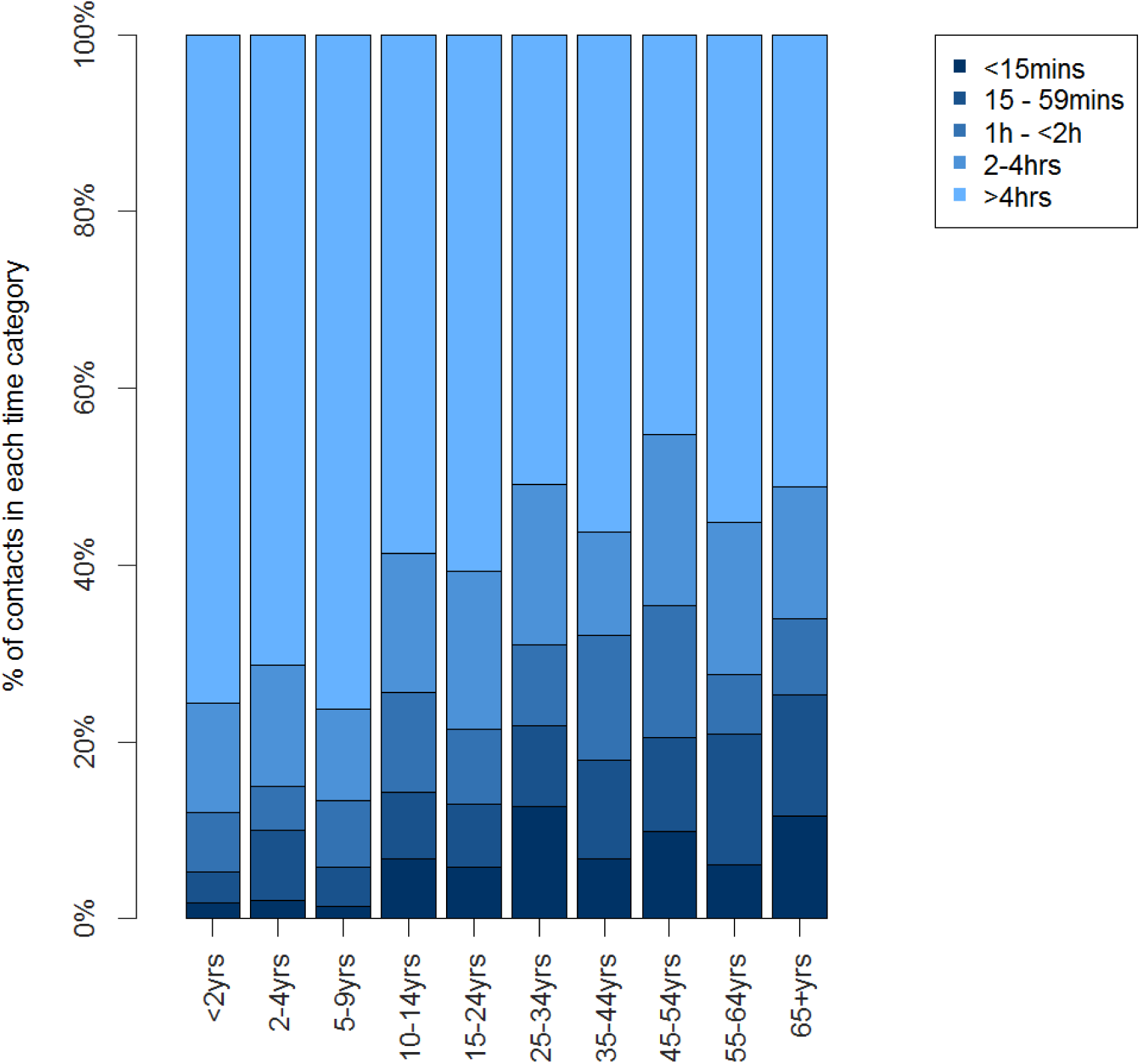
The Reported Duration of Contact By Age Group Among Study Participants, Sheema District, January – March 2014

**Figure S2.**
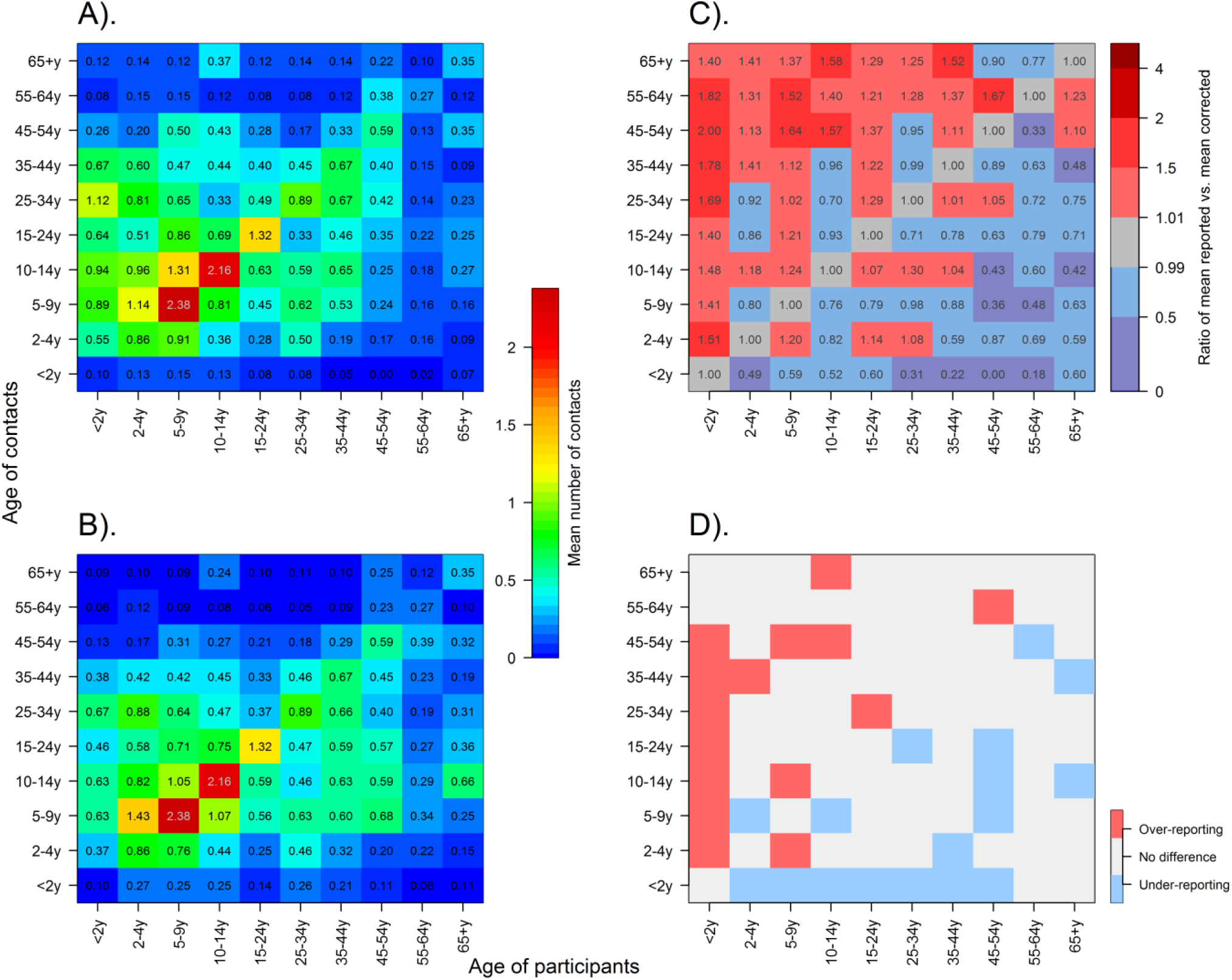
Reciprocity Correction of the Contact Matrices for All Contacts, Sheema District, January – March 2014. <u>Legend:</u> A) matrix for all reported contacts, not corrected. B) matrix for all reported contacts, corrected for reciprocity. C) Ratio of corrected over uncorrected matrices. Red cells illustrate where age-specific contacts were over-reported before correction, and blue cells under-reported. D) shows where participants significantly over-reported the number of age-specific contacts they had (upper 95% confidence bound) in red, significantly under-reported contacts in blue (lower 95% confidence bound), or where no significant adjustment was made (grey)

**Figure S3.**
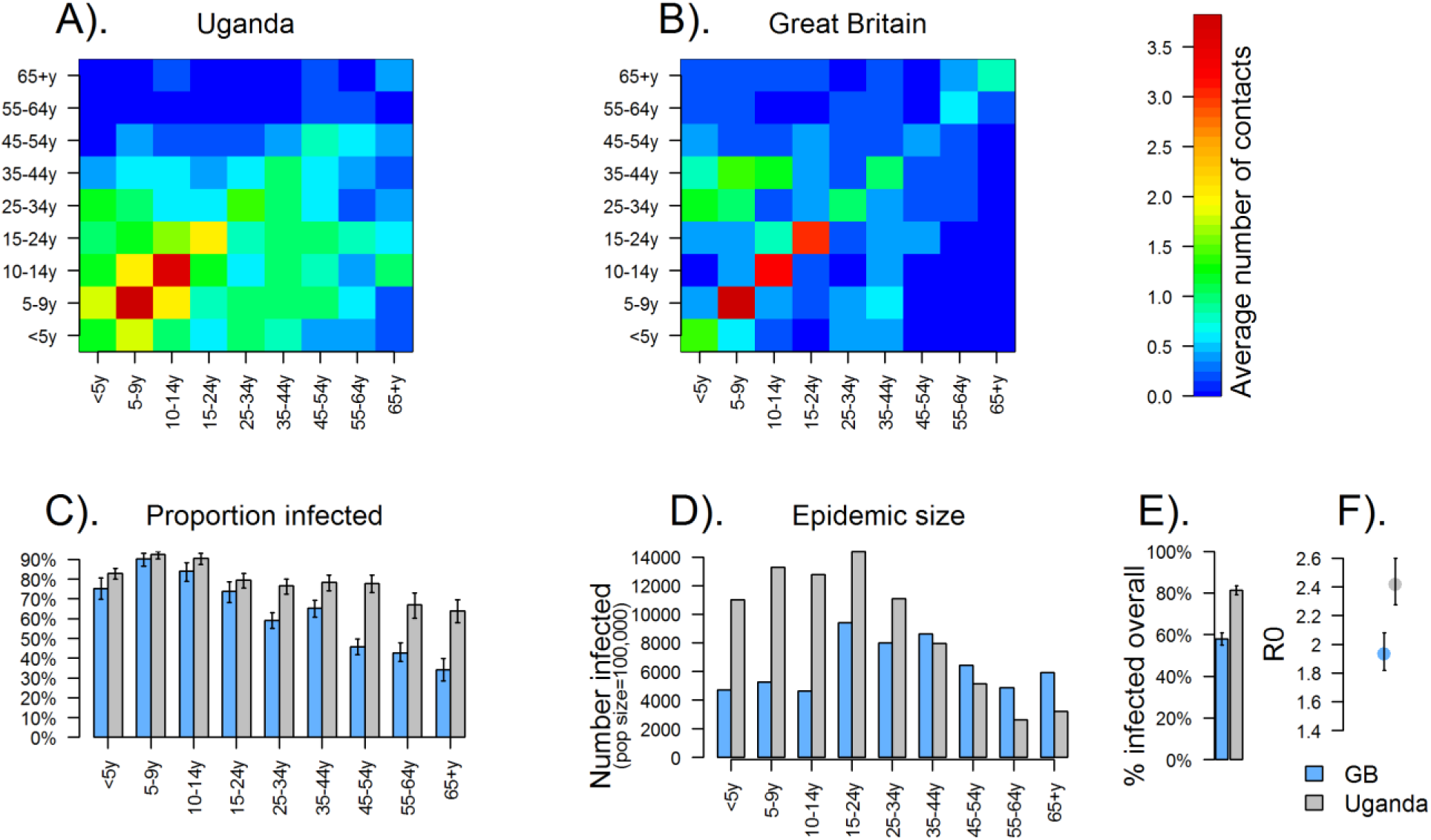
Epidemic Simulations Using Comparing Uganda and Great Britain, Assuming a 25% Underreporting Of Contacts In Uganda. <u>Legend:</u> A) Physical Contacts from Uganda, B) Physical Contacts from Great Britain, C) Proportion Infected by Age Group, D) Epidemic Size by Age group, E) Overall Proportion Infected, and F) the Basic Reproduction Number R0.

